# Is there an intrinsic relationship between LFP beta oscillation amplitude and firing rate of individual neurons in monkey motor cortex?

**DOI:** 10.1101/586727

**Authors:** Joachim Confais, Nicole Malfait, Thomas Brochier, Alexa Riehle, Bjørg Elisabeth Kilavik

## Abstract

It is a long-standing controversial issue whether an intrinsic relationship between the local field potential (LFP) beta oscillation amplitude and the spike rate of individual neurons in the motor cortex exists. Beta oscillations are prominent in motor cortical LFPs, and their relationship to the local neuronal spiking activity has been extensively studied. Many studies demonstrated that the spikes of individual neurons lock to the phase of LFP beta oscillations. However, the results concerning whether there is also an intrinsic relationship between the amplitude of LFP beta oscillations and the firing rate of individual neurons are contradictory. Some studies suggest a systematic mapping of spike rates onto LFP beta amplitude, and others find no systematic relationship. To resolve this controversy, we correlated the amplitude of LFP beta oscillations recorded in motor cortex of two male macaque monkeys with spike counts of individual neurons during visuomotor behavior, in two different manners. First, in an analysis termed *task-related correlation*, data obtained across all behavioral task epochs was included. These task-related correlations were frequently significant, and in majority of negative sign. Second, in an analysis termed *trial-by-trial correlation*, only data from a fixed pre-cue task epoch was included, and correlations were calculated across trials. Such trial-by-trial correlations were weak and rarely significant. We conclude that there is no intrinsic relationship between the firing rate of individual neurons and LFP beta oscillation amplitude in macaque motor cortex, beyond each of these signals being modulated by external factors such as the behavioral task.

**SIGNIFICANCE STATEMENT:** We addressed the long-standing controversial issue of whether there is an intrinsic relationship between the local field potential (LFP) beta oscillation amplitude and the spike rate of individual neurons in the motor cortex. In two complementary analyses of data from macaque monkeys, we first demonstrate that the unfolding behavioral task strongly affects both the LFP beta amplitude and the neuronal spike rate, creating task-related correlations between the two signals. However, when limiting the influence of the task, by restricting our analysis to a fixed task epoch, correlations between the two signals were largely eliminated. We conclude that there is no intrinsic relationship between the firing rate of individual neurons and LFP beta oscillation amplitude in motor cortex.

## INTRODUCTION

The properties of motor cortical local field potential (LFP) beta oscillations were the focus of many studies. They occur as bursts (Murthy and Fetz 1996; Donoghue et al. 1998; Feingold et al. 2015), typically lasting 100-500ms and generally not locked to external events. They are however related to the task (event-related), such that the probability of observing beta bursts changes across task epochs (e.g. Feingold et al. 2015). Soon after their first description (Berger 1929), human sensorimotor beta oscillations were linked to states of neuronal activity equilibrium (Jasper and Penfield 1949). Subsequently, periods of beta event-related synchronization (ERS) and desynchronization (ERD) were interpreted as reflecting deactivation and activation, respectively, of the sensorimotor cortex (Pfurtscheller et al. 1996; Pfurtscheller and Lopes da Silva 1999; Pfurtscheller 2001; Salenius et al. 1997; Neuper et al. 2006; Bechthold et al. 2018). This concept mainly springs from the robust observations of much reduced beta oscillation amplitude just before and during movements (Kilavik et al. 2013). The notion that motor cortical beta ERD/ERS indexes neuronal activation/deactivation (Neuper et al. 2006) might suggest that one should expect an inverse relationship between neuronal spike rates and beta amplitude.

Several studies addressed the relationship between macaque motor cortical LFP beta oscillations and the local spiking activity (e.g. Murthy and Fetz 1996; Donoghue et al. 1998; Baker et al. 1999; Denker et al. 2011; Canolty et al. 2012; Engelhard et al. 2013; Best et al. 2017; Rule et al. 2017, 2018; Riehle et al. 2018). Importantly, the pioneering studies by Murthy and Fetz (1996) and Donoghue et al. (1998) studied the relationship between LFP beta amplitude and neuronal spike rates. The first study found no modulations in the rate of neurons in relation to beta amplitude (Murthy and Fetz, 1996), whereas the other found some motor cortical locations with increased firing rates during increased oscillation amplitude, and others showing the opposite (Donoghue et al. 1998). Unfortunately, these contradictory results remain overlooked in more recent, relevant studies.

Canolty et al. (2012) studied in great detail the relationship between LFP beta oscillations and neuronal spiking in macaque motor cortex. They demonstrated several distinct dependencies between LFP beta amplitude and the firing rates of individual neurons, which they termed ‘amplitude-to-rate’ mapping. Some neurons exhibited a negative correlation and others a positive correlation with beta amplitude. Furthermore, the amplitude-to-rate mapping of individual neurons could be reversed across behavioral contexts (manual vs. brain control task). They concluded that the dependency of spike rates upon beta amplitude (internal factor) was conditioned upon the specific behavioral task (external factor). Womelsdorf et al. (2013) therefore suggested that by means of this amplitude-to-rate mapping, beta activity could mediate switches between sub-networks across task epochs and across tasks. This supposes an intrinsic relationship between beta amplitude and firing rate.

More recently, Rule et al. (2017) found no consistent relationship between LFP beta amplitude and spike rates, but they did not discuss the contradictory finding of Canolty et al. (2012). Indeed, differences in data analysis approaches might be the cause of the different conclusions of these two studies. Canolty et al. (2012) analyzed data by including all task epochs. Rule et al. (2017) restricted their analysis to steady-state movement preparation periods.

To resolve this controversial issue, we correlated macaque motor cortical LFP beta oscillation amplitude with neuronal spike counts obtained during visuomotor behavior (Kilavik et al. 2012; Confais et al. 2012). When analyzing data including all behavioral task epochs, correlations were frequently observed, confirming the results of Canolty et al. (2012). However, when restricting the analysis to the pre-cue epoch, and performing a trial-by-trial correlation analysis, significant correlations were rare, confirming the results of Rule et al. (2017). We conclude that there is no intrinsic relationship between the firing rate of individual neurons and LFP beta oscillation amplitude in motor cortex, beyond simple co-modulations driven by task events. Some preliminary results were presented in Kilavik and Riehle (2015).

## MATERIALS AND METHODS

We analyzed LFP signals and spiking data recorded simultaneously on multiple electrodes in motor cortex of two macaque monkeys during the performance of a visuomotor delayed center-out reaching task. We used previously obtained data, from which other results have been presented (Kilavik et al. 2010, 2012, 2014; Ponce-Alvarez et al. 2010; Confais et al. 2012). We have already shown that this dataset contains strong LFP oscillations in the beta range, which are systematically modulated in amplitude and peak frequency by the behavioral task (Kilavik et al. 2012). We have also reported on robust and specific modulations in neuronal spiking activity in relation to the behavioral task (Confais et al. 2012). The experimental data can be shared upon request.

### Animal preparation and data recording

Two adult male Rhesus monkeys (T and M, both 9kg) participated in this study. Care and treatment of the animals during all stages of the experiments conformed to the European and French Government Regulations applicable at the time the experiments were performed (86/609/EEC).

After learning an arm-reaching task (see below) the monkeys were prepared for multi-electrode recordings in the right hemisphere of the motor cortex, contra-lateral to the trained arm. The recording chamber locations above primary motor and dorsal pre-motor cortex were verified with T1-weighted MRI scans in both monkeys, and also with intra-cortical micro-stimulation in monkey M (see details in Kilavik et al. 2010). Across all included recording locations, the sampled regions spanned about 4 and 13mm diameter on the cortical surface in monkeys T and M, respectively (Kilavik et al. 2010), and were in majority arm/hand related.

A multi-electrode, computer-controlled microdrive (MT-EPS, AlphaOmega, Nazareth Illith, Israel) was used to transdurally insert up to four or eight (in monkey T and M, respectively) microelectrodes. The reference was common to all electrodes and positioned, typically together with the ground, on a metal screw on the saline-filled metallic recording chamber. In monkey T the electrodes were organized in a bundle in one common larger guide tube holding the individual electrode guides, with an inter-electrode distance <400µm (MT; AlphaOmega). However, since the electrodes were driven independently, their position in depth varied for each electrode. In monkey M, on some days electrodes were organized in a bundle as for monkey T and on others the electrodes were positioned independently within the chamber with separate guide tubes (Flex-MT; AlphaOmega), thus resulting in up to 13mm inter-electrode distance. The amplified raw signal (1 Hz – 10 kHz) was digitized and stored at 32 kHz. For the online extraction of single neuron activity, the amplified raw signal was hardware high-pass filtered at 300Hz to obtain the high-frequency signal, on which an online spike shape detection method was applied (MSD, AlphaOmega, Nazareth Illith, Israel), allowing isolation of up to three single neurons per electrode. The timing of each spike was then stored as TTLs at a temporal resolution of 32 kHz, down-sampled offline to 1 kHz before analysis. Offline spike sorting on the raw signals was additionally performed in Matlab (The MathWorks Inc., USA) by using Principal Component Analysis in the toolbox MClust (http://www.stat.washington.edu/mclust/) when the online spike sorting was considered as non-optimal. In parallel, the amplified raw signal was hardware low-pass filtered online at 250Hz to obtain the low-frequency LFP signal, which was stored with a temporal resolution of 1 kHz. Behavioral data were transmitted online to AlphaMap (AlphaOmega) from the CORTEX software (NIMH, http://dally.nimh.nih.gov), which was used to control the task.

### Behavioral task

We trained the two monkeys to make arm-reaching movements in 6 directions in the horizontal plane from a common center position, by holding a handle that was freely movable in the two-dimensional plane (Figure 1A). In some sessions, only 2 random-chosen opposite directions were used to reduce the session duration, concerning 21% and 39% of the analyzed sessions in monkey T and M, respectively. The monkeys had continuous feedback about hand (white cursor) and the 6 possible target positions (red outlines) on a vertical monitor in front of them.

**Figure 1:**
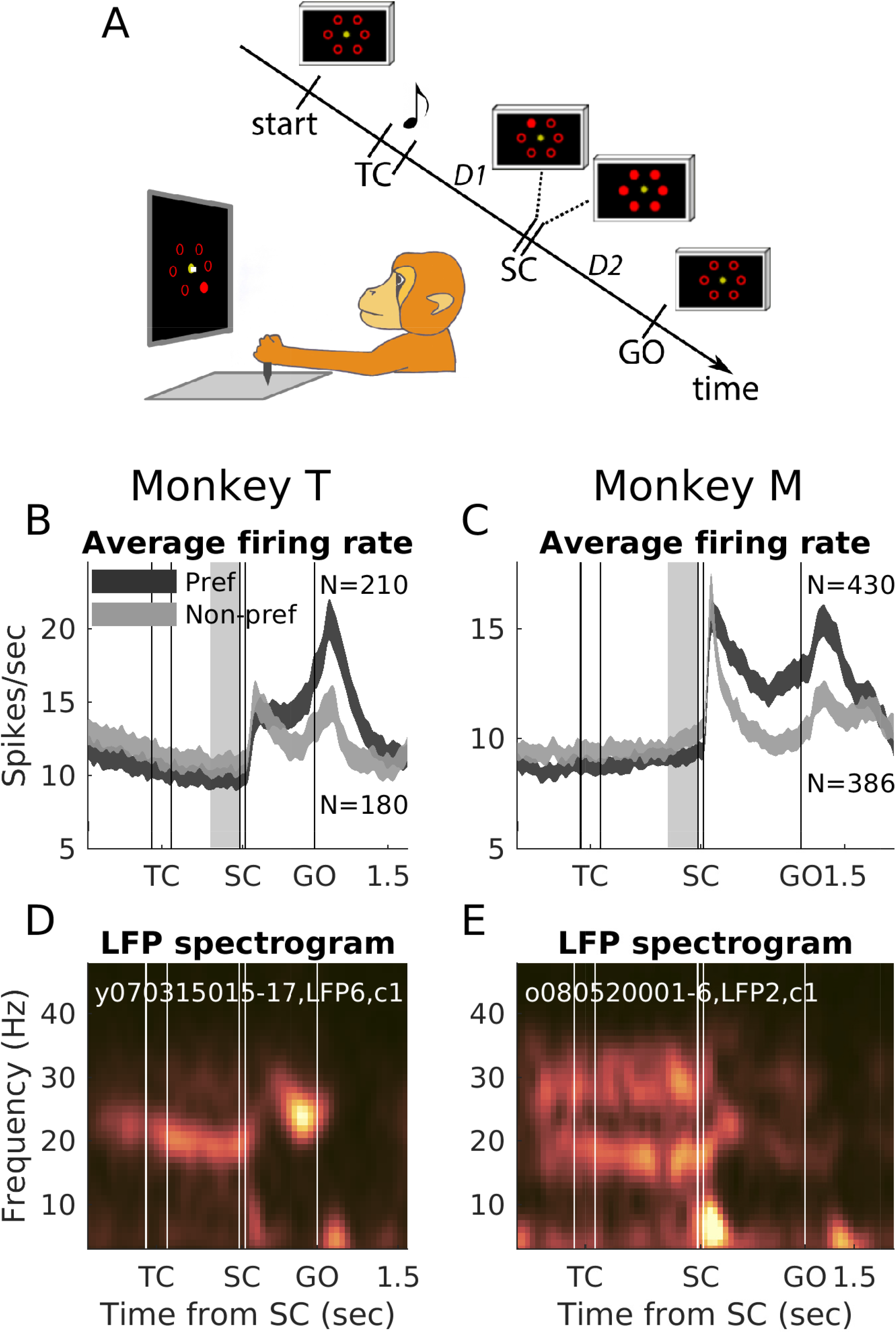
Behavioral paradigm, average neuronal rates and example LFP spectrograms. A: Behavioral paradigm. Left, drawing of the experimental apparatus showing the SC epoch (note the cursor on the central fixation dot). Right, Sequence of task events, not to scale. Start indicates the moment when the monkey brings the cursor to the center of the screen to initiate a new trial. The musical note indicates the presentation of a tone. Tone pitch differs according to delay duration. All screen-shots shown in the diagram stay on until the next one appears (cursor is not shown). TC, 200 ms; SC, 55 ms; D1, delay 1, D2, delay 2. Both delays have either short duration (700ms in monkey T and 1000ms in monkey M) or long duration (1500ms in monkey T and 2000ms in monkey M). There is also a 700ms delay between start and TC. B-C: Average rate for all neurons included in the task-related correlation analysis, for preferred (dark gray) and non-preferred (light gray) movement directions, in short delay trials for monkey T (left) and monkey M (right). The curves reflect the mean firing rate +/− SEM. Data between trial start and until 1000ms after the GO signal (as depicted) was included in the task-related correlation analysis. The epoch marked in light gray preceding SC was used for the trial-by-trial correlation analysis. The average rate for each SUA was smoothed with a Gaussian filter of length 50ms and sigma 20ms, before averaging. N in the plots reflects the number of included neurons. Note the reduced numbers of neurons for the non-preferred direction, caused by imposing an average minimal rate of 3Hz for each direction separately (see methods). This selection criterion also causes a somewhat higher population firing rate from the start of the trial for the non-preferred direction. Since per definition the rate is lower after SC for the non-preferred compared to the preferred direction, the somewhat fewer neurons for the non-preferred direction have slightly higher rate from the start of the trial. D-E: Spectrograms of one representative example LFP for each monkey, including all correct short delay trials in one condition (session, LFP number and condition indicated inside plots). Frequency is on the vertical axis and time along the horizontal axis. Warmer colors indicate increased power (a.u.) using a perceptually flat color-map (Crameri 2018), with color limits set to the minimum and maximum power values above 10Hz, separately for each monkey. To create the spectrograms, the LFPs were first high-pass filtered at 2Hz with a 4^th^ order Butterworth filter before the power spectral density (based on discrete Fourier transform) was calculated, at 1Hz frequency resolution. The averages across all trials were plotted at the center of each sliding window (300ms duration, 50ms shifts). The brief power-increases below 10Hz after SC and GO reflect visual and movement evoked potentials.

Two delays were presented successively in each trial. The two delays (D1 and D2) had the same fixed duration, either short or long. Their duration was instructed by an auditory cue just before D1 initiation, set from trial to trial in a pseudo-random fashion. Their durations were either 700 or 1500ms for monkey T, 1000 or 2000ms for monkey M. The monkey started each trial by moving the handle to the center (‘start’ in Figure 1A) and holding it there for 700ms until a temporal cue (TC) was presented. TC consisted of a 200ms long tone, its pitch indicating the delay duration, starting at the end of the tone (low pitch for short and high pitch for long delay duration). The delay that followed TC (D1) involved temporal attention processes (Confais et al. 2012), to perceive the spatial cue (SC) that was illuminated very briefly (55ms) at the end of the delay at one of the peripheral target position. To assure the temporal precision of SC illumination time and duration, light-emitting diodes (LEDs) were used, which were mounted in front of the computer screen in fixed positions at the center of the 6 peripheral red target outlines, on a transparent plate. SC was subsequently masked by the additional illumination of the 5 remaining LEDs, marking the start of D2. During D2 the movement direction indicated by the visual cue SC had to be memorized and prepared. All LEDs went off at the end of D2 (GO signal), indicating to the monkey to reach towards and hold (for 300ms) the correct peripheral target position. In summary, during D1 the monkey had to wait for SC, which was briefly presented at the end of the pre-cued time interval. D2 entailed visuomotor integration and movement preparation while waiting for the GO signal.

The reaction and movement times were computed online to reward the monkey after target hold, with a maximum allowance of 500ms for each. For data analysis, the reaction times were redefined offline using the arm trajectories. Trajectories were measured in x and y vectors at 1ms resolution. The mean of each x and y vector during the 500ms before GO in each trial was used as the movement’s starting position. The moment when reaching a 2mm deviation, minus a fixed latency of 35ms (average movement duration from the starting position to the threshold), was determined as movement onset. From each of the two vectors (x and y), the shortest time was defined as RT. These values were controlled by visual inspection of single trial trajectories (see Kilavik et al. 2010).

### Data selection and analysis

While the monkeys performed the reaching task we recorded neuronal activity from motor cortex. We recorded 90 sessions in 37 days in monkey T and 151 sessions in 73 days in monkey M. Consecutive sessions in the same day were made after lowering further the electrodes to sample new neurons. This provided a total of 287 and 759 individual recording sites in monkeys T and M, respectively. A site is here defined as the conjunction of a specific chamber coordinate of the electrode entry and the cortical depth. After site elimination due to lack of sufficiently recorded trials, or large recording artifacts affecting either the lower (LFP) or higher (spiking activity) frequencies, 127 and 358 sites remained for further analysis, from 66 and 135 individual sessions, for monkeys T and M, respectively. These essentially constitute the conjunction between the LFP datasets studied in Kilavik et al. (2012) and the single neuron datasets studied in Confais et al. (2012).

All analyses were conducted offline by using Matlab (The MathWorks, Inc.). We studied a low beta band that was strong in both animals. In addition, in monkey M who also had a marked beta band at higher frequency (see Kilavik et al. 2012 and example in Fig. 1E), the analysis was repeated for this band. We first band-pass filtered the LFP around the average peak beta frequency for each band with a zero-phase 4^th^ order Butterworth filter. In monkey T the LFPs were filtered between 22+/-5Hz to capture the dominant low beta band across the entire trial (see example in Figure 1D and averages across all LFPs in Kilavik et al. 2012). For monkey M, to capture the low and high beta bands across the entire trial the LFPs were filtered at 19+/-5Hz and 32+/-5Hz, respectively (see Figure 1E and Kilavik et al. 2012). After filtering, beta oscillation amplitude was estimated from the analytical filtered LFP, as the envelope of the signal from the Hilbert transform.

From the online and offline spike sorting, typically 1 to 3 neurons were available on each electrode. For the correlation analyses between LFP beta amplitude and neuronal firing rate, beta-neuron pairs were constructed using signals from different simultaneously recorded sites. This choice was guided by findings demonstrating the possibility of spike contamination of LFP signals recorded on the same electrode, also for the lower LFP frequency ranges studied here (Zanos et al. 2011; Waldert et al. 2013). From the 127 and 358 acceptable sites, 320 and 671 beta-neuron pairs were constructed in monkeys T and M, respectively. Each neuron was paired with only one LFP. As describe in the data recording details above, we used two different electrode microdrives with different inter-electrode spacing. For all pairs in monkey T, and 61% (408/671) of pairs in monkey M, the co-recorded site used for LFP beta was less than 400µm away from the spiking site in chamber coordinates (but at different cortical depths). The remaining 39% (263/671) of pairs in monkey M were constructed with sites typically 1-6mm apart, with a few sites up to 11 mm apart. Whenever multiple co-recorded sites were available, the site selection for LFP beta was mainly driven by LFP signal quality. Different LFPs recorded up to 1mm apart in motor cortex typically show very similar modulations in beta amplitude on a trial-by-trial basis (see Kilavik et al. 2012).

In these pairs, some trials with obvious artifacts (mainly due to teeth grinding or static electricity) detected by visual inspection, were excluded from further analysis (less than 5% of all trials). After trial elimination, and considering the variable duration for which the monkeys were willing to work in different behavioral sessions, the analyzed beta-neuron pairs contained at least 10 correct trials in each movement direction, although typically 20 or more correct trials were available per direction. The average numbers of correct trials in each direction (in short or long delay trials) across pairs were 23+/-5 (mean +/− standard deviation) for monkey T and 20+/-5 for monkey M. The average numbers of total short (long) delay trials for each pair were 117+/-36 (117+/-37) for monkey T and 93+/-36 (90+/-36) for monkey M.

### Task-related correlations between LFP beta amplitude and neuronal spike counts

We here define *task-related correlation* as the correlation between two brain signals calculated across several diverse task epochs, such that the concurrent modulations in the brain signals related to the unfolding task events and related behavior can be expected to influence the amount of correlation observed between them. The task-related correlation was calculated between LFP beta oscillation amplitude and neuronal spike counts for each beta-neuron pair. Data recorded in all epochs between the trial start (initial central touch) and until 1000ms after the GO signal was included (as displayed in Figures 1B-C and 2A-B), analyzed separately for short and long delay trials. Across the included sessions, the average reaction times in short (long) delay trials were 161 (206) ms in monkey T and 232 (255) ms in monkey M, and the average movement times were 303 (296) ms in monkey T and 297 (303) ms in monkey M (see also Kilavik et al. 2010). Thus, average reaction and movement times were both shorter than their maximally allowed durations of 500ms each, so that the analysis typically also includes most of the required 300ms target-hold time.

**Figure 2:**
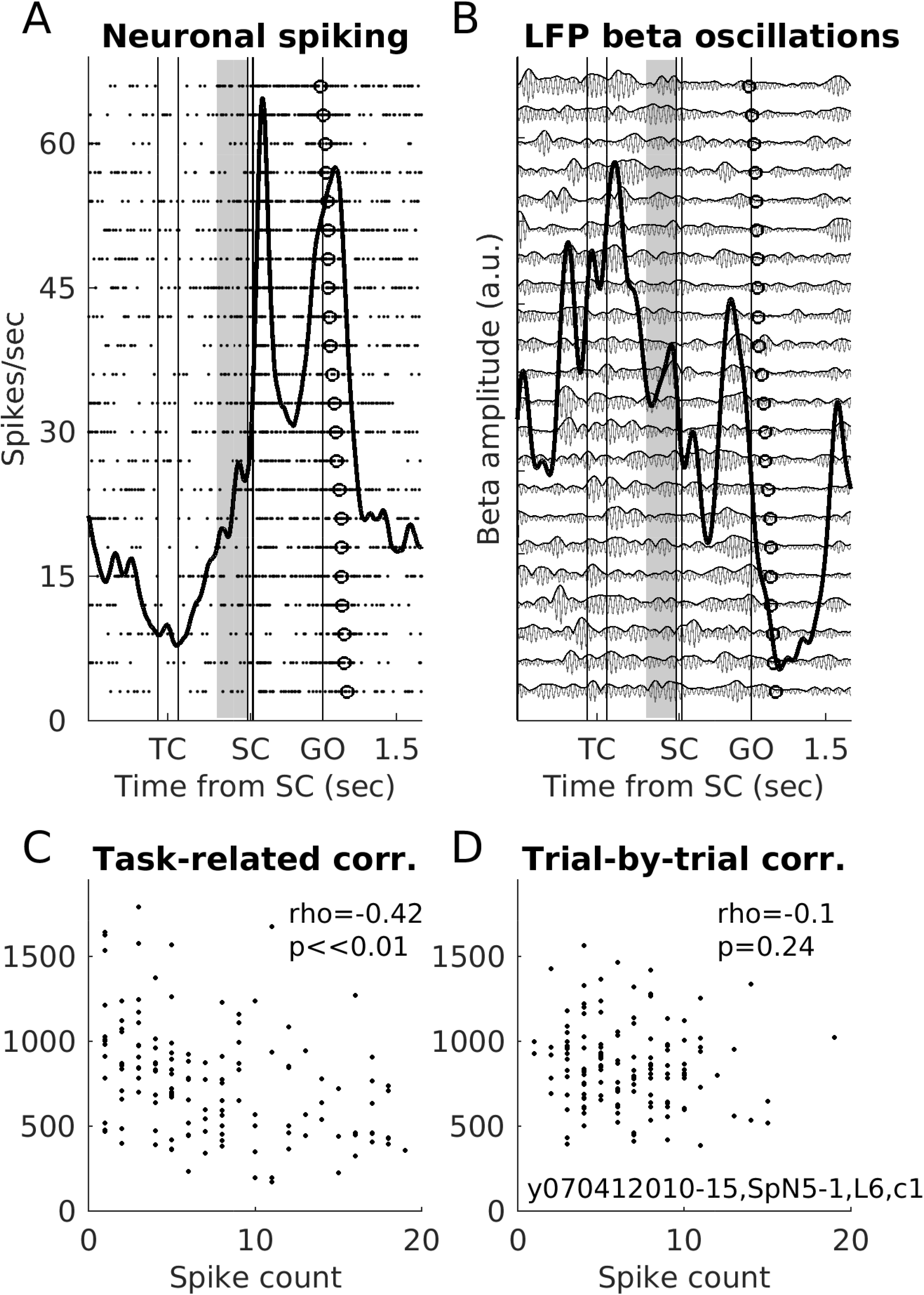
Example beta-neuron pair. A: Raster plot and peri-stimulus time histogram (PSTH) of one example neuron in its preferred movement direction, in short delay trials (example from monkey T; session, neuron and LFP ID, and condition indicated inside plot D). In the raster plot, each dot is an action potential and each row a trial, ordered according to reaction times (open circles; shortest on top). The thick black line represents the neuronal activity averaged across all the shown trials (PSTH; smoothed with a Gaussian filter of length 100ms and sigma 50ms). The epoch marked in gray preceding SC (also in B) was used for trial-by-trial correlation analysis shown in D (that also included all short delay trials for all the other movement directions). B: LFP from another co-recorded electrode, filtered for the beta range (22+/-5Hz; light gray curves), shown for the same individual trials as the raster plot for the neuron in A. Darker gray curves show the instantaneous beta oscillation amplitude, which was estimated from the analytical filtered LFP as the envelope of the signal from the Hilbert transform. The thick black line indicates the average beta amplitude across all shown trials (smoothed with a Gaussian filter of length 100ms and sigma 50ms). C: This pair’s task-related correlation, for short delay trials in the preferred direction. The results from selecting only as many windows as number of trials (n=135) is shown. Each dot corresponds to one 300ms window, with combined values of beta amplitude and spike counts. The Spearman’s rho was −0.42, a highly significant negative correlation. D: This pair’s trial-by-trial correlation, for short delay trials (n=135). Each dot corresponds to the beta amplitude and spike counts for one trial, in the 300ms pre-SC window marked in gray in A-B. The correlation was not significant.

The beta-neuron correlations were calculated separately for the preferred and non-preferred (opposite) movement direction for the neuron in each pair, where preferred direction was taken as the one with maximal trial-averaged spike rate any time after the presentation of SC up to trial end. This was done to evaluate whether the task-related correlation with LFP beta amplitude depended on the involvement of the neuron in coding for the cued movement.

The single trial data in these two directions was cut in 300ms non-overlapping consecutive windows. The window duration of 300ms was in part chosen based on the typical duration of beta bursts in our dataset (200-500ms), see example in Figure 2B; see also Murthy and Fetz 1992). Note that recent literature suggests that in some contexts beta bursts can be of much shorter duration than seen in our dataset (e.g. Feingold et al. 2015; Sherman et al. 2016; Lundqvist et al 2016). Since our 300ms windows were aligned to the task timing, i.e. signal occurrences, and beta bursts do not have a fixed temporal relationship with such external events, some windows will overlap with a beta burst, while others will fall in a period with low beta amplitude, and some will partly overlap with a beta burst.

Secondly, window duration of 300ms was considered to be a minimal duration needed for meaningful (non-zero) spike counts in a majority of individual windows. However, we additionally restricted our analysis to the subsets of beta-neuron pairs for which the average firing rate of the neuron, across all 300ms windows, was above 3 Hz. The numbers of analyzed pairs thus varied slightly for short and long delay trials and for preferred and non-preferred movement directions, as detailed in Table 1 (see also Figure 1B-C).

**Table 1.** Summary of results from correlation analyses. Proportions (numbers and percentages) of beta-neuron pairs with significant task-related and trial-by-trial correlations, presented separately for monkey T, and monkey M low (LO) and high (HI) beta bands. The proportions of significant correlations with negative sign are specified. Short and long delay trials and neuronal preferred and non-preferred movement directions are presented separately. For the task-related correlation analysis, we present results obtained when using all available windows, and with corrected (reduced) number of windows, to have as many windows as trials.

| Analysis type | Data selection | Monkey T |  | Monkey M LO |  | Monkey M HI |  |
| --- | --- | --- | --- | --- | --- | --- | --- |
|  |  | Proportions of significant correlations | Proportions of negative correlations | Proportions of significant correlations | Proportions of negative correlations | Proportions of significant correlations | Proportions of negative correlations |
| Task-related correlation<br>All available windows | Short delay<br>Pref. dir. | 104/210<br>(50%) | 84/104 (81%) | 177/430<br>(41%) | 133/177<br>(75%) | 205/430<br>(48%) | 174/205<br>(85%) |
|  | Short delay<br>Non-pref. dir. | 69/180 (38%) | 33/69 (48%) | 139/386<br>(36%) | 83/139 (60%) | 167/386<br>(43%) | 104/167<br>(62%) |
|  | Long delay<br>Pref. dir. | 88/204 (43%) | 72/88 (82%) | 175/412<br>(42%) | 140/175<br>(80%) | 202/412<br>(49%) | 165/202<br>(82%) |
|  | Long delay<br>Non-pref. dir. | 74/181 (41%) | 36/74 (49%) | 138/374<br>(37%) | 87/138 (63%) | 153/374<br>(41%) | 97/153 (63%) |
| Task-related correlation<br>Corrected number of windows | Short delay<br>Pref. dir. | 65/209 (31%) | 57/65 (88%) | 77/422 (18%) | 63/77 (82%) | 113/422<br>(27%) | 93/113 (82%) |
|  | Short delay<br>Non-pref. dir. | 44/177 (25%) | 26/44 (59%) | 61/384 (16%) | 38/61 (62%) | 84/384 (22%) | 49/84 (58%) |
|  | Long delay<br>Pref. dir. | 56/199 (28%) | 45/56 (80%) | 62/413 (15%) | 50/62 (81%) | 76/413 (18%) | 70/76 (92%) |
|  | Long delay<br>Non-pref. dir. | 35/186 (19%) | 18/35 (51%) | 45/376 (12%) | 22/45 (49%) | 60/376 (16%) | 34/60 (57%) |
| Trial-by-trial correlation | Short delay | 7/178 (3.9%) | 3/7 | 21/374<br>(5.6%) | 5/21 | 17/374<br>(4.5%) | 8/17 |
|  | Long delay | 6/176 (3.4%) | 4/6 | 18/378<br>(4.8%) | 9/18 | 14/378<br>(3.7%) | 6/14 |

This trial cutting provided 11 (16) non-overlapping 300ms windows in monkey T and 13 (19) in monkey M, for short (long) delay trials. The total number of windows accumulated across trials varied because of variable number of correctly performed trials across sessions. The average numbers of overall available windows for all trials in the same (preferred or non-preferred) movement direction in short (long) delay trials were 259+/-64 (373+/-96) for monkey T and 283+/-77 (400+/-113) for monkey M. The average beta amplitude (Hilbert envelope; see Figure 2B) and the spike counts in each 300ms window (providing one value per signal type in each window) was then used to calculate the beta-neuron task-related correlation, quantified with the Spearman’s rank order correlation (Spearman’s rho). Correlations with p<0.01 were considered significant, but the complete distributions of rho values across the populations of beta-neuron pairs are always presented, to allow appreciating the magnitude of the different types of correlations.

The analysis approach just described resembles as closely as possible for our dataset the approach used by Canolty et al. (2012). They concatenated LFP and spike data across several recording sessions from implanted multi-electrode arrays, providing between 58-410 minutes of continuous data, including all task epochs. However, in our dataset, there were on average more than twice as many windows available for this task-related correlation analysis approach compared to the number of trials available for the trial-by-trial correlation analysis described in the next section (averages of 117 trials in both short and long for monkey T and 93 and 90 trials in short and long, respectively, for monkey M; see above). This difference may pose problems in comparing the results due to sample size affecting the statistical power. To permit a more direct comparison between the task-related correlation analysis and the trial-by-trial correlation, the analysis was repeated after selecting from the total available windows a subset equaling the number of short (or long) delay trials for each beta-neuron pair. As far as possible, this selection was done such that every second window was excluded. The selection of every second window was repeated if there were still too many windows. Finally this selection was complemented with additional (previously excluded) windows if needed, to arrive at the correct number of windows. In the Results section we describe task-related correlations using both analyses (including all, or corrected number of windows); summarized in Table 1. However, in figures we only present results using corrected number of windows (example pair in Figure 2C and population distributions in Figure 3A-B).

**Figure 3:**
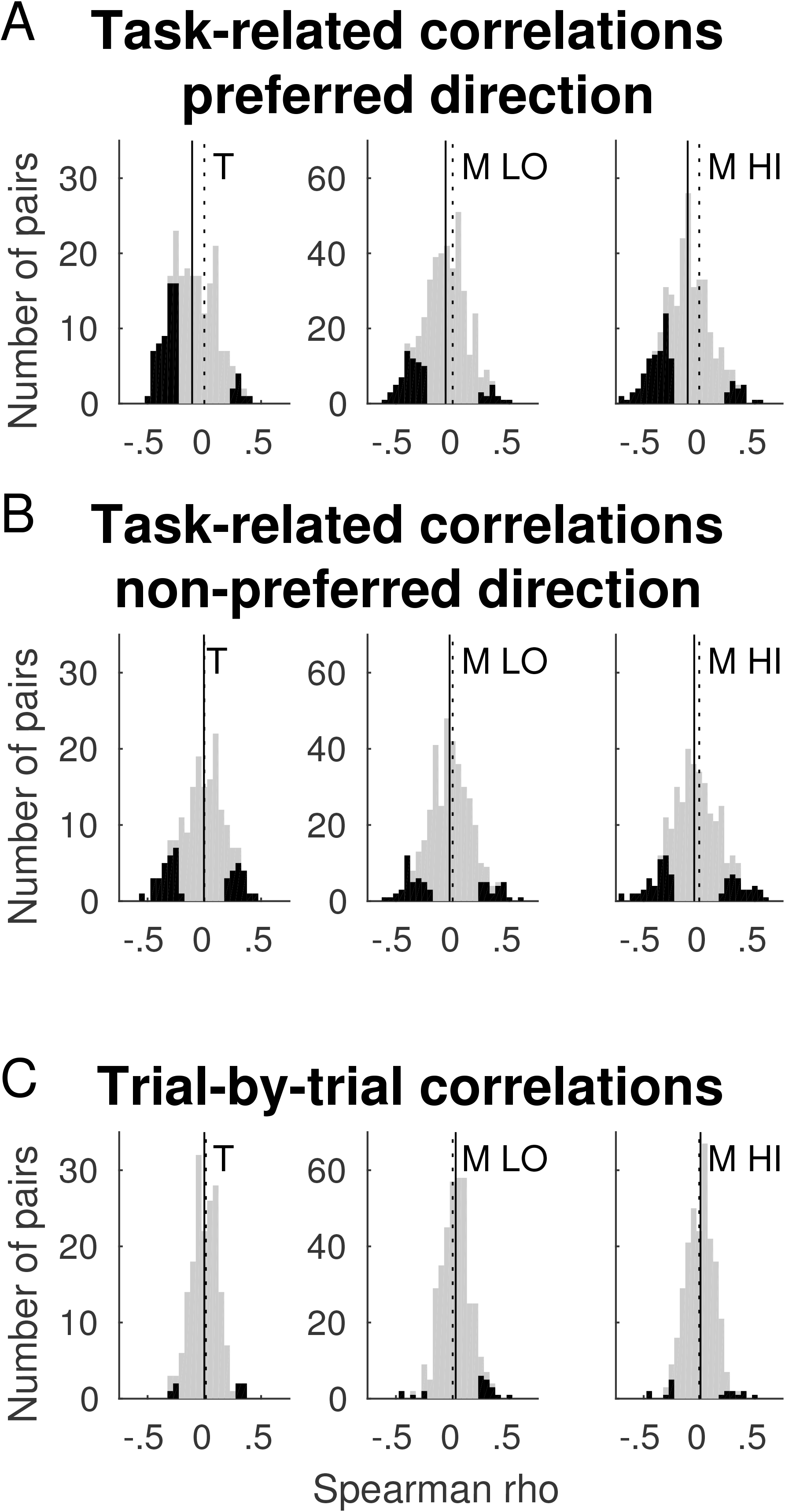
Task-related and trial-by-trial correlations. A: Complete distributions of Spearman’s rho values for task-related correlations in neuronal preferred movement direction in short delay trials, for all pairs in gray and overlaid in black for the significant pairs (p<0.01), for monkey T (left) and monkey M low beta band (LO; middle) and high beta band (HI; right). Dotted lines mark zero. Solid lines mark the medians of the complete (gray) distributions, which were significantly shifted to the left (negative correlations; Wilcoxon signed rank test on Fisher’s z-transformed rho values; p<<0.01 for all datasets). B: Distributions of task-related correlations for neuronal non-preferred movement direction in short delay trials. For monkey M low and high bands, the distributions were significantly shifted to the left (negative correlations; p<0.01), while for monkey T the distribution was centered on zero (p=0.66). Further details as in A. C: Distributions of trial-by-trial correlations in short delay trials. The distributions were centered on zero for monkey T (p=0.46) and monkey M for the high band (p=.22), and only slightly shifted to the right (positive correlations, p<0.01) for monkey M for the low band. Further details as in A-B.

### Trial-by-trial correlations between LFP beta amplitude and neuronal spike counts

For each beta-neuron pair the *trial-by-trial correlation* between LFP beta oscillation amplitude and neuronal spike counts was calculated in a 300ms epoch immediately preceding SC, across all trials, separately for short and long delay trials (gray epoch in Figure 2A-B). Since our 300ms window was aligned to SC, and beta bursts do not have a fixed temporal relationship with the external events (see introduction, and example in Fig. 2B), as for the task-related correlation analysis explained above, on some trials the window overlaps with a beta burst, while in some trials it will fall in a period with lower beta amplitude, and in some trials it will partly overlap in time with a beta burst.

We choose this restricted task moment by considering it to be the epoch in which the monkey’s behavioral state was most likely to be similar across all trials within each delay duration condition. Specifically, the monkey maintained a stable arm position on the central target and was awaiting the presentation of a visual cue. Notably, this epoch started between 1.3-2.6 seconds after the monkey had moved his hand cursor into the central target to start a new trial, and 0.4-1.7 seconds after the end of the presentation of the auditory temporal cue (TC off) providing information about delay duration. In this epoch the movement direction was still unknown, so all directions can be grouped in the analysis, while analyzing short and long delay trials separately. Significant trial-by-trial beta-neuron correlations in this epoch may be mainly related to modulations of internal (anticipatory) processes, thereby reflecting any intrinsic beta-spike relationship, independent of external factors related to the task such as the processing of external visual or auditory sensory cues or overt movements. This analysis is more closely comparable to the analysis performed by Rule et al. (2017), in which they restricted their analysis to delays considered to entail steady-state movement preparation. However they compared epochs inside and outside of beta bursts, thus at varying moments in their 1 second duration delays considered. As we already described for the current dataset, the neuronal spike rates modulated significantly from the start to the end of the delay (D1) preceding SC, some neurons systematically increasing and others decreasing their rate between TC and SC (Confais et al. 2012). Thus, in our case, this delay cannot be considered as steady-state in its entirety. To avoid these systematic, task-related modulations in neuronal firing rates influencing our analysis, the window was restricted to the final 300ms prior to the pre-indicated moment of SC onset.

As for the task-related correlation analysis, the analysis included only the subsets of beta-neuron pairs for which the average firing rate of the neuron in the pre-SC epoch was above 3 Hz. The LFPs were filtered to capture the main beta frequency band(s) for each animal as described above. The average beta amplitude (Hilbert envelope) and the spike counts in each trial in this 300ms epoch for analysis (providing one value per signal type per trial window) were then correlated across trials. The trial-by-trial correlation for each beta-neuron pair was then quantified as the Spearman’s rank order correlation (see example in Figure 2D; results across all datasets in Table 1 and Figure 3C), as described above for the task-related correlation analysis.

### Variability in spike counts and beta amplitude

To determine to which degree the correlation analyses results were dependent upon the level of signal variability, we estimated the variability of the spike counts and beta amplitude across analyses windows. This was done for each beta-neuron pair and for data entering the task-related correlation in neuronal preferred direction and for the trial-by-trial correlation, separately for short and long delay trials. We calculated the coefficient of variation (CV; standard deviation divided by the mean). The CV of each type of signal was then correlated with the Spearman’s rho values from the beta amplitude – neuronal spike count correlations across pairs.

### Phase-locking of neuronal spiking to LFP beta phase

To confirm that the LFP beta oscillations were at least partially of local origin, we verified that a substantial proportion of the neurons significant locked their spiking activity to the LFP beta phase. The proportion of neurons with a significant phase-locking to beta oscillations was quantified in a 300ms duration pre-SC epoch, separately for short and long delay trials. We focused on this particular task epoch since one of the analyses of correlations between beta-amplitude and neuronal spike counts was done on this same epoch (the trial-by-trial correlation). To ensure a reliable statistical analysis, only neurons with at least 50 spikes in this 300ms epoch, accumulated across all trials, were included. This restricted the analysis to a subset of 229 (226) of the 320 pairs in monkey T and 441 (448) of the 671 pairs in monkey M, for short (long) delay trials. Beta phase was extracted from the Hilbert transformation of the beta-filtered LFP, and the phase at each spike time was determined.

To quantify the phase locking, we first used Rayleigh’s test of non-uniformity of circular data (CircStat Matlab toolbox; Berens 2009). To determine whether the locking was significant for individual neurons, a trial-shuffling method was used. Beta oscillations are typically not phase-reset by external events (but see Reimer and Hatsopoulos 2010), and the analyzed pre-SC epoch was sufficiently long after the previous external event (0.4-1.7s after TC off), such that any phase-resetting effects should have minimal effect in this epoch. This makes trial-shuffling an efficient method for obtaining a ‘baseline’ measure of phase locking, destroying the temporal relationship between the two signals, while preserving their individual properties such as rhythmicity.

In the trial-shuffling analysis, 1000 repetitions of the phase-locking analysis (Rayleigh’s test; in the same 300ms pre-SC epoch) was done while randomly combining beta phases and spike times from different trials. If the original analysis yielded a larger z-statistic value from the Rayleigh’s test than 990/1000 (equivalent to p<0.01) of the trial-shuffled analyses, the phase-locking of the neuron was considered to be significant.

## RESULTS

The aim of this study was to determine to which degree there is an intrinsic relationship between the amplitude of LFP beta oscillations and firing rates (spike counts) of individual neurons in the motor cortex. We correlated motor cortical LFP beta amplitude and neuronal spike counts measured in short windows either along the trial including several different task epochs (*task-related correlation*) or within a fixed task epoch, but across trials (*trial-by-trial correlation*). We start with a general description of the average task-related modulation in firing rate of the included neurons, as well as the typical task-related modulations of LFP beta amplitude.

### Modulations in neuronal firing rates and LFP beta amplitude during task performance

The monkeys performed a visuomotor arm-reaching task (Figure 1A), while we recorded neuronal activity from motor cortex. Figure 1B-C shows the average firing rates of all neurons included in this study, separated for neuronal preferred and non-preferred movement direction. At the population level there was a phasic increase in rate for both the preferred and non-preferred directions following the spatial cue (SC). The population rate then decreased during the preparatory delay between SC and GO, but remained above the pre-SC level in particular for the preferred direction, before increasing again towards and during movement execution after GO.

Example LFP spectrograms for each monkey are shown in Figure 1D-E. These examples are representative when it comes to the average beta power and frequency across task epochs in these datasets, as we already described in detail in Kilavik et al. (2012). Notably monkey T had one dominant beta band, which varied in average frequency between 19-25Hz across task epochs. Monkey M had two dominant beta bands, a low band modulating between 17-21Hz and a high band modulating between 29-34Hz (Kilavik et al. 2012). For both monkeys and both bands, beta power decreased after SC and during movement execution after GO. Note that even if these trial-averaged spectrograms suggest a prolonged increase in beta amplitude during the delays, as can be seen in the example LFP in Figure 2B in reality beta oscillations occur in individual bursts of different duration, amplitude and exact timing across trials (see also Feingold et al. 2015; Sherman et al. 2016; Lundqvist et al 2016).

### Task-related correlations between LFP beta amplitude and neuronal spike counts are prominent

We calculated task-related correlations between LFP beta oscillation amplitude and neuronal spike counts along the trial including different task epochs for the 320 and 671 beta-neuron pairs in monkeys T and M, respectively. An example pair with significant task-related correlation is shown in Figure 2C. This particular pair showed a negative correlation between beta amplitude and neuronal firing rate. The overall percentages of significant correlations, for both monkeys and beta bands, in short and long delay trials and in the neuronal preferred and non-preferred movement directions are summarized in Table 1.

Task-related correlations using all available windows were prominent and frequently significant for both monkeys, and for both beta bands in monkey M. The complete distributions of Spearman’s rho values were rather broad and significantly shifted towards negative values (Wilcoxon signed rank test on Fisher’s z-transformed rho values; p<<0.01; distributions for the results using all windows not shown).

The task-related correlations for the preferred direction were statistically significant (p<0.01) in 41-50% of pairs (across both monkeys and bands, short and long delay trials; Table 1). A combination of negative and positive significant correlations was observed, as also described by Canolty et al. (2012). However, the large majority of the significant correlations for neuronal preferred directions were negative (75 to 85% across both monkeys and bands, short and long delay trials). This dominance of negative correlations is possibly due to the systematic decreases in beta amplitude following the visual spatial cue (SC) and during movement execution (see Figure 1D-E; Kilavik et al. 2012, 2013), which occurs more or less concurrently with phasic increases in firing rates in a majority of neurons in their preferred direction (see Figure 1B-C and Confais et al. 2012).

In order to evaluate whether the task-related correlation with LFP beta amplitude depended on the involvement of the neuron in coding for the cued movement, we also analyzed neuronal non-preferred movement direction. Here, 36-43% of pairs had significant task-related correlations, across both monkeys and bands, short and long delay trials (Table 1). However, the proportions of these significant correlations being negative were smaller than for the preferred direction (48-63% across both monkeys and bands, short and long delay trials). After a brief phasic increase in rate following the spatial cue, which at the population level is similar in preferred and non-preferred movement directions (see Figure 1), the neurons discharge less in the non-preferred compared to the preferred movement direction, and some neurons discharge less than their pre-cue rate. This could be expected to lead to larger proportions of neurons having a positive correlation with beta amplitude for their non-preferred direction, as beta amplitude also drops after the cue and during movement execution. The proportions of significant negative and positive correlations were very similar for short and long delay trials (Table 1), as would be expected if the movement directional preferences of the neurons were the major cause for the sign of the beta-neuron task-related correlations.

When comparing pairs with significant correlations in both the preferred and the non-preferred directions, of the pairs significant in both directions, only a small fraction changed correlation sign, mainly from negative in preferred to positive in non-preferred (e.g. in short delay trials 4/44 in monkey T, 2/95 and 4/108 in monkey M low and high bands; only 1 pair changed correlation sign from positive to negative, for monkey M low beta band). Thus, the different proportions of significant negative correlations for preferred vs. non-preferred directions mainly stem from pairs being significantly correlated in only one of the directions. The changes in the sign of task-related correlations between preferred and non-preferred movement directions at the population level can therefore not be interpreted as a ‘remapping’ in the relationship to beta amplitude for individual neurons, in the way described by Canolty et al. (2012) when switching between their manual and brain control tasks.

In order to have comparable statistical power as for the trial-by-trial correlation analysis, for which the results will be described in the next section, the task-related correlation analysis was also done by selecting only as many windows as there were available trials for the trial-by-trial correlation analysis for each individual pair. These are therefore also the results shown in the distribution plots in Figure 3A-B, for short delay trials. This correction of number of windows reduced the overall proportions of pairs with significant task-related correlations (15-31% for neuronal preferred movement direction, across both monkeys and bands, short and long delay trials), which is not surprising since we reduce statistical power by reducing the sample sizes. However, the main results of broad distributions of rho values (Figure 3A), and a majority of significant correlations being negative for the preferred direction (80-88% across both monkeys and bands, short and long delay trials; Table 1) remained similar. All the distributions of Spearman’s rho values were significantly shifted towards negative values (Figure 3A). The distributions of rho values remained broad also for the non-preferred direction using the corrected number of windows (Figure 3B), but as for the preceding analysis, the proportions of the significant correlations (12-25% across both monkeys and bands, short and long delay trials) having negative sign decreased compared to the preferred direction (49-62% across both monkeys and bands, short and long delay trials; Table 1). Furthermore, the distributions were only significantly shifted away from zero for monkey M.

There was a gradual decrease in the proportions of pairs with significant task-related correlations going from preferred direction in short delay trials to non-preferred direction in long delay trials (see Table 1). This might be due to more gradual modulations across task epochs of both beta burst probability and spike counts for longer delays, scaled to delay duration (discussed in Kilavik et al. 2014), in addition to some neurons having shallower modulations across task epochs for their non-preferred direction (Figure 1B-C).

In general these results are in agreement with the findings by Canolty et al. (2012), that they interpreted as a ‘beta-to-rate mapping’, with a specific relationship between the firing rates of individual neurons and the amplitude of beta oscillations.

### Trial-by-trial correlations between LFP beta amplitude and neuronal spike counts are rare

Figure 2D shows that in the selected example beta-neuron pair, LFP beta amplitude and neuronal spike count did not correlate trial-by-trial in the pre-SC epoch. This was indeed representative of the populations. Only 3.4-5.6% of the pairs had a significant correlation (across both monkeys and bands, short and long delay trials), with similar proportions of negative and positive correlations (see Table 1). Figure 3C shows the distributions of Spearman’s rho values for the pre-SC trial-by-trial correlation analysis in short delay trials for the three datasets. The distributions were narrower than for the task-related correlations, and only significantly shifted away from zero for the low beta band in Monkey M in short delay trials, not in long delay trials.

These very weak and rarely significant trial-by-trial correlations are in line with the results in Rule et al. (2017), where they describe inconsistent differences in firing rates for low and high beta amplitude events in their steady-state preparatory period analysis.

### No influence of signal variability on beta-neuron correlations

In order to estimate to which degree significant task-related or trial-by-trial correlations were associated with neuron pairs having large signal variability, we quantified the variability (CV) of the neuronal spike counts and beta amplitudes across analyses windows.

For the spike counts, the population distributions of CV magnitudes were slightly smaller (two-sample t-tests; p<0.01) across the windows entering the trial-by-trial correlation than across the task-related windows, except for monkey M for short delay trials (p=0.016). However, across beta-neuron pairs, spike-count CV correlated neither with the strength of task-related nor trial-by-trial correlations (Spearman’s rank order correlation; p>0.01 for all comparisons). In other words, those neurons with large spike count variability were not more likely to be correlated with beta amplitude.

Beta amplitude was much less variable across trials in the pre-SC epoch than across windows included in the task-related correlation analysis (two-sample t-tests; p<<0.01 for all comparisons). This was to be expected due to the large fluctuations in beta burst probabilities particularly comparing delays to the post-cue and movement epochs. Still, as can be appreciated in the example in Figure 2D, the mean beta amplitude could still triple on some trials compared to others in the pre-SC epoch. As for spike counts, beta amplitude CV correlated neither with beta-neuron trial-by-trial nor task-related correlation strength (Spearman’s rank order correlation; p>0.01). The only significant association was for an increased beta amplitude variability for pairs with higher task-related correlations for long delay trials for the low beta band in monkey M (p=0.008).

### Neurons lock their spikes to LFP beta oscillation phase

The LFP is prone to containing a combination of signals generated by local and distant sources (e.g. Kajikawa and Schroeder 2011). When wanting to study the relationship between the LFP beta oscillation amplitude and local spiking activity it is essential to verify the likewise local origin of the beta oscillations. A significant phase-locking of the spiking activity of the local neuronal population reveal locking of the neurons to synchronized synaptic inputs (of local or distant origin), in turn leading to local postsynaptic currents that contribute to generating the LFP (Pesaran et al. 2018). As a control analysis, we therefore confirmed that the spiking activity of a significant proportion of the neurons locked to the phase of the LFP beta oscillations. This control was specifically done in the pre-SC epoch, where only very few neurons with a trial-by-trial spike count modulation in relation to beta amplitude were found, as described in the previous section.

Overall, in the analyzed pre-SC epoch, 37.6% and 40.7% of the neurons locked significantly their spiking activity to the beta phase of the LFP in monkey T in short and long delay trials, respectively. 11.1% and 12.2% of the neurons locked significantly to the low and high beta bands, respectively, in monkey M in short delay trials, and 8.0% and 14.3% were phase-locked to the low and high band in long delay trials. In monkey M, only 2.0% in short delay trials and 2.7% in long delay trials of the neurons locked significantly their spikes to both the low and the high bands in this task epoch, such that overall 21.3% in short delay trials and 19.6% in long delay trials of the neurons locked their spikes to either the low, high or both LFP beta bands in this monkey. The clear phase-locking found for many neurons in this dataset made us conclude that the observed LFP beta bands were at least partly locally generated, justifying the correlation analyses between beta amplitude and neuronal firing rate. Finally, there was no systematic difference in locking prevalence of the few neurons with, compared to without, a significant trial-by-trial correlation of spike count with beta amplitude in the pre-SC task epoch.

## DISCUSSION

To reconcile two apparently contradictory results about the relationship between beta amplitude and neuronal firing rate, we here performed systematic quantifications of correlations between macaque motor cortical LFP beta amplitude and spike counts in individual neurons during a visuomotor task, in two different manners. First, in the analysis called *task-related correlation*, analogous to the approach by Canolty et al. (2012), data obtained across all behavioral task epochs were included. Such task-related correlations were frequent and in majority negative. Second, in the analysis called *trial-by-trial correlation*, analogous to the approach by Rule et al. (2017), only data from a fixed pre-cue epoch were included, and the trial-by-trial correlation of beta amplitude and spike counts was calculated. We found such trial-by-trial correlations to be very rare. We conclude that there is no intrinsic dependency between neuronal spike count and beta amplitude, beyond both types of signals being modulated by external factors such as the behavioral task.

### Disparate literature evidence for an intrinsic relationship between motor cortical beta amplitude and neuronal firing rates

The question of whether modulations in beta amplitude are related to modulations in the activation level of local neurons was already examined more than 20 years ago. In a behavioral context in which macaques made reaching movements to a Klüver board, Murthy and Fetz (1996) found no difference in average firing rates of individual neurons inside and outside beta bursts (20-40Hz) in motor cortex. However, they found a decrease in the variability of firing rates of individual neurons during and just after burst events, compared to just before bursts. They also noted that many neurons were phase-locked to the high-amplitude beta oscillations, which might be the main source of this decreased firing rate variability. Donoghue et al. (1998) analyzed LFPs and neuronal discharge (individual neurons and multi-units) during tasks involving finger or arm movements. One group of multi-units ‘overlapped’ with LFP oscillations (20-60Hz), increasing their discharge in epochs of increased oscillation amplitude. Another, ‘mixed’ group mainly decreased their discharge during increased beta oscillation amplitude, but also showed some overlap. They noted that the consistent patterns for each recorded site suggested the two signals (LFP amplitude and neuronal rate) be mechanistically linked.

These two rather contradictory early studies (Murthy and Fetz 1996; Donoghue et al. 1998) cannot be directly compared, since their methods have significant differences (using the spiking activity of single or multi units; considering different LFP frequency ranges; differences in behavioral tasks). Furthermore, they were unfortunately not cited by subsequent literature addressing the same question. More recently, Canolty and colleagues (2012) presented a rigorous analysis of the ‘cross-level coupling’ between spikes and beta oscillations, and described an ‘amplitude-to-rate mapping’. Some neurons exhibited a strong negative correlation and others a strong positive correlation with beta amplitude, and this mapping could change across tasks (manual or brain control tasks; Canolty et al. 2012). The notion of an amplitude-to-rate mapping supposes an intrinsic relationship between beta amplitude and firing rate, and might be interpreted such that beta activity indexes switches between sub-networks across different task epochs, and different tasks (Womelsdorf et al. 2013).

Rule et al. (2017) also addressed the same question, finding no consistent relationship between beta amplitude and spike rates when restricting their analysis to steady-state preparation periods. Noteworthy, Engelhard et al. (2013) trained macaques to increase motor cortical 30-40Hz LFP oscillation power and spike synchrony, and found no systematic modulation in neuronal firing rates when comparing low and high LFP power periods.

### Reconciling these findings – no intrinsic relationship beyond co-modulations driven by task events

A direct comparison of the two recent studies (Canolty et al. 2012; Rule et al. 2017) suggests that their discrepant conclusions might be due to different analysis approaches, either including data from all trial epochs, or restricting their analysis to steady-state preparatory periods, respectively. The two ways in which the data were analyzed in this study, quantifying both *task-related* and *trial-by-trial* correlations in the same dataset, resemble the approaches used by Canolty et al. (2012) and Rule et al. (2017), respectively. Indeed, we confirm the results of Canolty et al. (2012) when including many different task epochs, and we confirm the results of Rule et al. (2017) when restricting our analysis to a fixed pre-cue epoch, using data across trials. Whereas Rule et al. (2017) compared firing rates inside and outside of beta bursts across a one second delay period, we used a fixed pre-SC window that was not always aligned to the beta bursts. Still, the same conclusion is reached. Importantly, we found no systematic relationship between variability in spike counts or beta amplitude and the strength of task-related or trial-by-trial correlations across pairs. Thus, the handful of beta-neuron pairs with a significant trial-by-trial correlation did not correspond to those with particularly high variability in spike counts or beta amplitude. Importantly, this suggests that per see the smaller signal variability in the pre-SC epoch is unlikely causing the much smaller proportions of pairs with significant trial-by-trial correlations.

The impact of movement initiation upon beta-neuron ‘task-related’ correlations was recently demonstrated by Khanna and Carmena (2017), who only analyzed beta amplitude and neuronal firing rates in the epoch surrounding movement onset, confirming and extending the findings of Canolty et al. (2012). Our results therefore reconcile the disparate results from these recent papers, and possibly also the results obtained by Murthy and Fetz (1996) and Donoghue et al. (1998). Interestingly, we obtained very similar results for both beta bands in monkey M. Thus no clear distinction can be made concerning potential functional roles of each band in this study, beyond the conclusion that there is no intrinsic relationship between beta oscillation amplitude and spike counts of individual neurons for any of the two bands.

To evaluate to which degree this task-related correlation with LFP beta amplitude depended on the involvement of the neuron in coding for the upcoming movement, we also analyzed non-preferred movement direction. The proportions of significantly correlated pairs were comparable for preferred and non-preferred movement directions. Furthermore, the different proportions of significant negative correlations resulted from pairs being significant in only one direction. The shift in population task-related correlation distributions towards the center for the non-preferred direction can therefore not be interpreted as a ‘remapping’ in the relationship to beta amplitude for individual neurons, in the way described by Canolty et al. (2012) across tasks. It however favors the idea that these beta-neuron correlations simply reflect to which degree the two signals are co-modulated by the behavioral task.

### Phase-locking of spikes to LFP beta oscillations

The lack of an intrinsic relationship between LFP beta amplitude and neuronal activation level (rate) does not exclude other relationships between beta oscillations and neuronal spiking activity. Indeed, as we demonstrate in this dataset, confirming several previous studies (Murthy and Fetz 1996; Donoghue et al. 1998; Baker et al. 1999; Denker et al. 2011; Canolty et al. 2012; Engelhard et al. 2013; Riehle et al. 2018) there is significant locking of spike times to LFP beta oscillation phase for many neurons in motor cortex. Such phase locking may result in rhythmic synchronization among populations of neurons thereby increasing their concerted impact on post-synaptic targets without necessary increasing their spike rates (Destexhe and Paré 1999; Azouz and Gray 2000).

### No need for several processes underlying motor cortical beta amplitude modulations

Rule et al. (2017) pointed out that beta amplitude decreases at movement onset, roughly when neurons in motor cortex are generally mostly active (see also Khanna and Carmena 2017; Best et al. 2017). This observation was in contradiction to the lack of a systematic relationship between beta amplitude and firing rates in their main analysis. They therefore proposed that two different processes govern motor cortical beta amplitude variability. One underlies the beta amplitude decrease around movement onset and is linked to large modulations in spiking rates. Another underlies the transient beta bursts during steady-state delays, lacking overt movements and decoupled from modulations in spiking activity.

Instead, we propose that there is no intrinsic relationship between LFP beta amplitude and neuronal firing rates. Thus, the significant task-related correlations observed in this study, as well as the beta-to-rate mapping described in Canolty et al. (2012) is rather a reflection of the beta amplitude (burst probability) and firing rates (spiking probability) both being modulated by the task events, however independently from each other. This implies no need for different processes underlying modulations of beta bursts in steady-state situations and for the suppression of beta bursts during movement execution (as well as after visual cues, see Kilavik et al. 2013; Zaepffel et al. 2013), as proposed by Rule et al. (2017). Even if the underlying generating mechanism might remain the same, this does not exclude potentially different functional roles for beta oscillation bursts occurring during cue anticipation, during movement preparation or post-movement (Kilavik et al. 2013; Torrecillos et al. 2015).

As a concluding remark, beyond understanding the mechanistic role of beta oscillations as observable in the intra-cortical LFP, this issue is highly relevant for studies in closely related fields using human participants. An extensive body of literature inquires the relationship between beta oscillations and task behavior, aiming at mechanistic understanding using non-invasive techniques in the human. It is crucial that we understand the relationship between these oscillations and the underlying spiking activity of individual neurons, across different levels of temporal precision, ranging from precise phase-locking to slower amplitude modulations.

## Acknowledgements

### Funding

This work was supported by the Agence Nationale de la Recherche [grant number ANR-NEUR-05-045-1]; Ministère de l’enseignement supérieur et de la recherche (PhD grant to J.C.). The authors wish to thank Adrian Ponce-Alvarez for participating in animal training and data recording; Ivan Balansard and Marc Martin for animal care; and Joel Baurberg, Alain De Moya, and Xavier Degiovanni for technical assistance. The current address of JC is Cynbiose, 69280 Marcy l’Étoile, France.

